# Sostdc1 regulates natural killer cell maturation and cytotoxicity

**DOI:** 10.1101/387225

**Authors:** Alberto J. Millan, Sonny R. Elizaldi, Eric M. Lee, Jeffrey O. Aceves, Deepa Murugesh, Gabriela G. Loots, Jennifer O. Manilay

**Affiliations:** University of California - Merced, Department of Molecular Cell Biology, School of Natural Sciences, Merced, CA, 95343, USA; Lawrence Livermore National Laboratories, Physical and Life Sciences Directorate, Livermore, CA 94550, USA.

## Abstract

Natural killer (NK) cells are specialized lymphocytes with the innate ability to eliminate virally infected and cancerous cells, but the mechanisms that control NK cell development and cytotoxicity are incompletely understood. We identified novel roles for Sclerostin domain containing-1 (*Sostdc1*) in NK cell development and function. *Sostdc1*-knockout (*Sostdc1*^-/-^) mice display a progressive accumulation of transitional NK cells (CD27^+^CD11b^+^, tNK) with age, indicating a partial developmental block. The Ly49 repertoire on NK cells in *Sostdc1*^-/-^ mice is also changed. Lower frequencies of *Sostdc1*^-/-^ splenic tNKs express inhibitory Ly49G2 receptors, but higher frequencies express activating Ly49H and Ly49D receptors. However, the frequencies of Ly49I^+^, G2^+^, H^+^ and D^+^ populations were universally decreased at the most mature (CD27^-^CD11b^+^, mNK) stage. We hypothesized that the Ly49 repertoire in *Sostdc1*^-/-^ mice would correlate with NK killing ability, and observed that *Sostdc1^-/-^* NK cells are hyporesponsive against MHC-I-deficient cell targets *in vitro* and *in vivo*, despite higher CD107a surface levels and similar IFNγ expression to controls. Consistent with Sostdc1’s known role in the regulation of Wnt signaling, high levels of Wnt coactivators *Tcf7* and *Lef1* were observed in *Sostdc1*^-/-^ NK cells. Expression of the NK development gene *Id2* was decreased in *Sostdc1^-/-^* iNK and tNK cells, but we observed no changes in *Eomes* and *Tbx21* expression. Reciprocal bone marrow transplant experiments showed that *Sostdc1* regulates NK cell maturation and expression of Ly49 receptors in a cell-extrinsic fashion from both non-hematopoietic and hematopoietic sources. Taken together, these data support a role for *Sostdc1* in the regulation of NK cell maturation, and NK cell cytotoxicity, and identify potential NK cell niches.

**Summary of Results:**

- *Sostdc1*^-/-^ mice display a partial block between the tNK and mNK developmental stages.
- *Sostdc1* influences the Ly49 receptor repertoire on NK cells.
- NK cells in *Sostdc1*^-/-^ mice display impaired ability to kill *β*_2_m^-/-^ target cells.
- *Sostdc1*^-/-^ NK cell subsets express high levels of Wnt coactivators *Tcf7* and *Lef1*.
- *Id2* expression is decreased in iNK and tNK cells in the absence of *Sostdc1*.
- Bone marrow transplantation experiments demonstrate cell-extrinsic regulation of NK cell maturation by Sostdc1 in both non-hematopoietic (stromal) and hematopoietic cells.

## Background and Rationale

Natural Killer (NK) cells are innate lymphocytes that are important for early immune defense against tumors and virally infected cells. Since the initial discovery of NK cells in 1975, studies from many groups have identified NK cell receptors that are involved in self/non-self recognition, NK cell precursors and stages of maturation, cytokines and transcription factors that are critical for NK cell development and function, and evidence for NK cell immune memory^1–9^. Despite over 40 years of NK cell history, the molecular and cellular mechanisms that drive and integrate these processes is still unclear. In particular, how the microenvironment regulates NK cell maturation and function is still an area of ongoing investigation.

Conventional NK cells develop in the bone marrow (BM) from hematopoietic stem cells (HSCs) following a well-established sequence of maturational stages and egress to the peripheral organs to fully mature and function^10,11,12^. NK cell maturation **(Figure 1A)** originates with the immature NK (CD27^+^CD11b^-^, iNK) cells, which progresses to the transitional NK cell stage (CD27^+^CD11b^+^, tNK), then to the final mature stage (CD27^-^CD11b^+^, mNK)^13–16^. As NK cells progress through these stages, they lose proliferative and cytokine-producing capability, but gain cytotoxic ability against target cells^12,17,18^. Although the BM microenvironment is critical for NK cell development, how the peripheral microenvironment regulates NK cell maturation and cytotoxicity is incompletely understood and requires further investigation.

**Figure 1.**
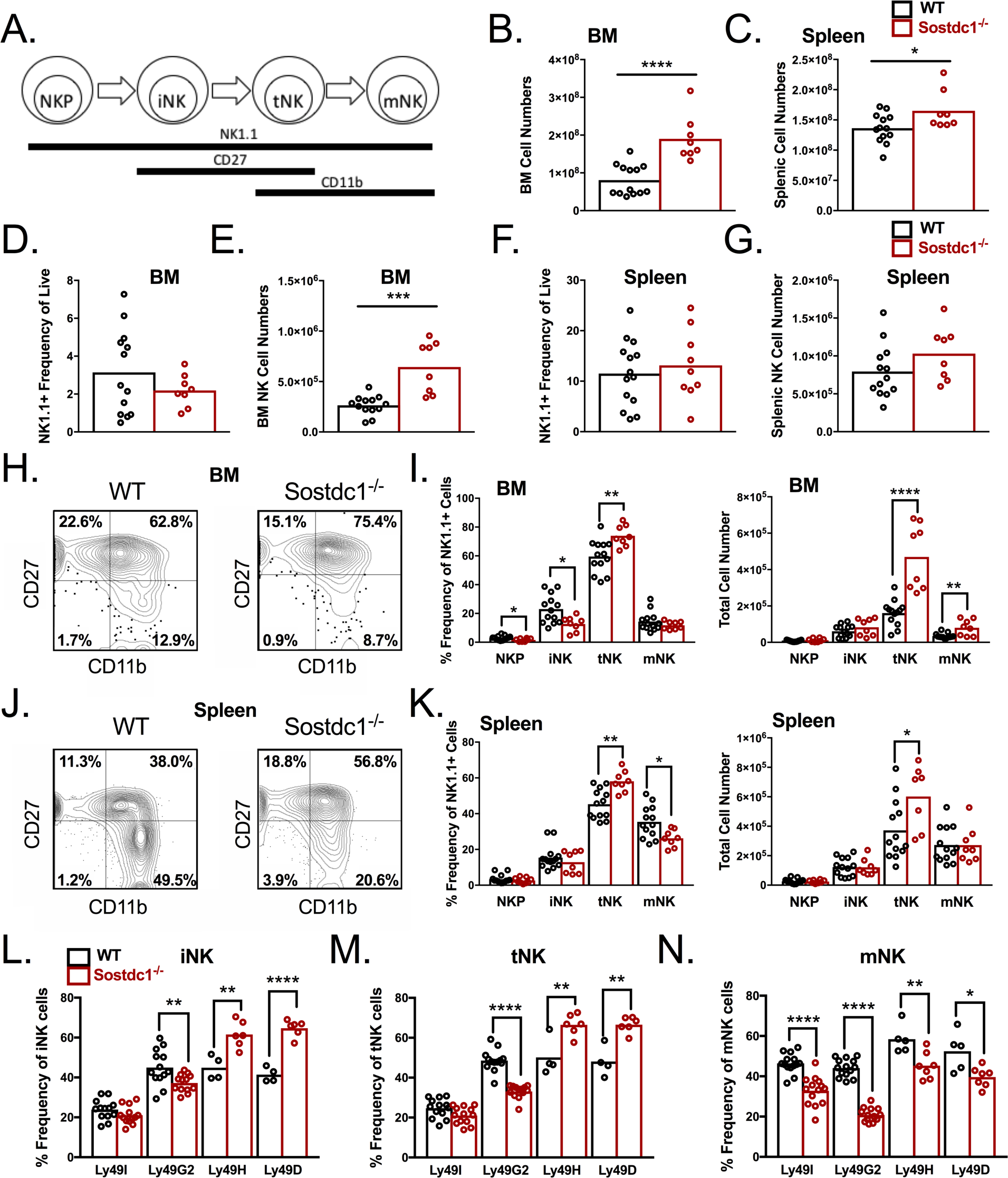
Delayed NK cell development and altered Ly49 repertoire in *Sostdc1^-/-^* mice. A) Schematic diagram of NK cell development: NKP: NK progenitor (CD27^-^ CD11b^-^), iNK: immature NK (CD27^+^ CD11b^-^), tNK: transitional NK (CD27^+^ CD11b^+^), mNK: mature NK (CD27^-^ CD11b^+^); B) Total cellularity of bone marrow (BM) in the femurs and tibiae and C) spleen of wild-type (WT) and *Sostdc1^-/-^* (KO) mice; D and F) Frequencies; and E and G) Absolute numbers of NK1.1^+^ cells in BM and spleen; H) Representative flow cytometry plots showing NK cell stages in BM; I) Summary of frequencies and absolute numbers of NKP, iNK, tNK and mNK cells in BM; J) Representative flow cytometry plots showing NK cell stages in spleen; K) Summary of frequencies and absolute numbers of NKP, iNK, tNK and mNK cells in spleen; L-N) distribution of splenic NK cells expressing Ly49I, Ly49G2, Ly49H and Ly49D on iNK, tNK and mNK cells. Asterisks indicate statistically significant differences between means, as determined by Student’s t-test. *p<0.05, **p<0.01, ****p<0.0001. Each point represents a single mouse.

Sclerostin domain containing-1 (*Sostdc1*), also known as Wise, Ectodin, Usag-1, and Sost-like, has been studied in the context of tooth development, kidney disease, hair follicle formation, and bone fracture^19–25^. Sostdc1 can function as an antagonist of both Bone Morphogenetic Protein (BMP) and canonical Wnt signaling pathways^20,21,23^. Sostdc1 expression is highly expressed in skin, brain, and intestine, as well as in skeletal muscles, kidney, lungs, and vasculature^20,22,23^. Most recently, we found it also to be expressed in the bone periosteum and mesenchymal stem cells (MSCs) to support bone formation and fracture remodeling^26^. Here, we reveal Sostdc1’s cell-extrinsic roles in the regulation of NK cell maturation, Ly49 receptor expression and cytotoxic function in the BM and spleen.

## Results

### Sostdc1^-/-^ mice display a partial block at the tNK stage

Our previous studies demonstrated that femurs of *Sostdc1*^-/-^ mice display a 21 percent increase in BM cavity volume compared to WT controls^26^. Consistent with this, the total BM cellularity of *Sostdc1*^-/-^ bones was increased (**Figure 1B**). *Sostdc1*^-/-^ mice also displayed higher total splenic cell numbers (**Figure 1C**). The increased cellularity suggested that the *Sostdc1*^-/-^ BM and spleen microenvironments may be altered, and that immune cell development may also be affected by the loss of *Sostdc1*. To test this, we performed flow cytometry (FCM) and observed no differences in frequencies or absolute numbers of CD19^+^ B lymphocytes, CD3^+^ T lymphocytes, and CD11b^+^ Gr1^+^ granulocytes (data not shown). However, the frequency of CD11b^+^ Gr1^-^ cells in the *Sostdc1^-/-^* spleen was reduced. To determine if the CD11b^+^ Gr1^-^ cells were monocytes or NK cells, we performed more detailed analysis with anti-NK1.1. Total NK (Live, CD3^-^,CD19-,Gr1^-^,NK1.1^+^) frequencies were not affected, but total NK cell numbers were increased only in the BM of *Sostdc1^-/-^* mice (**Figure 1D-1G**). We then investigated if lack of *Sostdc1* affected NK cell maturation (**Figure 1A**), and discovered that *Sostdc1*^-/-^ mice exhibit a partial block between the tNK (NK1.1^+^ CD11b^+^ CD27^+^) and mNK (NK1.1^+^ CD11b^+^ CD27^-^) cell stages in both the BM and spleen, as demonstrated by the increase in frequency and number of tNK cells in the BM (**Figures 1H and 1I**) and spleen (**Figures 1J and 1K**), and the decreased frequency of mNKs in the spleen. These data indicated that *Sostdc1* is required for full developmental progression from the tNK to the mNK cell stages.

### Absence of Sostdc1 alters the Ly49 receptor repertoire on NK cells

We also examined if NK cells in *Sostdc1^-/-^* mice expressed different levels and distributions of inhibitory (Ly49I and Ly49G2) and activating (Ly49D and Ly49H) Ly49 receptors. FCM analysis of the Ly49 repertoire on iNKs, tNKs, and mNKs in *Sostdc1*^-/-^ mice revealed decreased frequencies of Ly49G2^+^ cells at all NK cell stages in the BM (**Supplemental Figure 1A-1C**) and spleen (**Figure 1L-1N**). In contrast, frequencies of Ly49H^+^ iNK and tNK cells in the *Sostdc1^-/-^* BM and spleen were higher than controls, but the frequencies of Ly49H^+^ mNK cells were reduced in both tissues (although only statistically significant in the spleen). Similarly, frequencies of Ly49D^+^ in iNK and tNK cells were increased and reduced amongst the mNKs in the *Sostdc1^-/-^* BM and spleen (**Figure 1L-1N and Supplemental Figure 1A-1C**). The frequencies of Ly49I^+^ in iNK and tNK cells were similar to controls, but frequencies of Ly49I^+^ mNK cells were decreased in *Sostdc1*^-/-^ mice (**Figure 1N and Supplemental Figure 1C**). The median fluorescent intensity (MFI) of staining for Ly49G2 and Ly49H was reduced on *Sostdc1*^-/-^ BM mNK cells only, indicating a relatively minor effect of Sostdc1 on cell surface Ly49 receptor expression levels (**Supplemental Figure 1D-1K**).

Since it is theorized that NK cell activity is governed by the combined set of Ly49 receptors expressed on a given NK cell, we further compared the frequencies of WT and *Sostdc1^-/-^* NK cells that express different combinations of Ly49 receptors^27^ (Supplemental Figure 2). Higher frequencies of iNK and tNK cells expressing more activating than inhibitory receptors (i.e. “activating repertoires”) were observed in *Sostdc1^-/-^* mice (**Supplemental Figure 2D and 2E)**. However, lower frequencies of mNK cells with activating repertoires were observed (**Supplemental Figure 2F)**. Taken together, these data show the lack of *Sostdc1* influences the Ly49 receptor repertoire.

**Figure 2.**
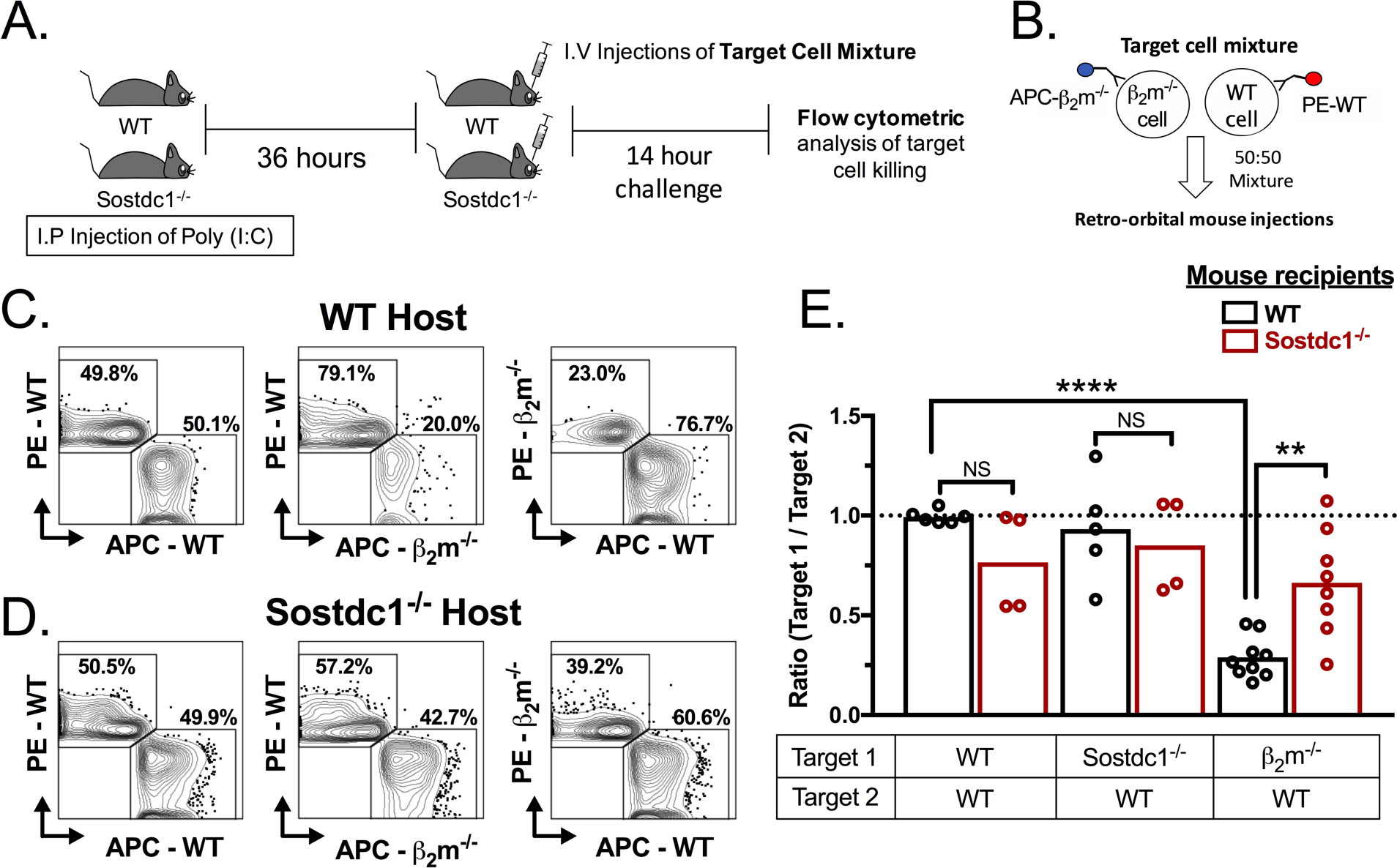
*Sostdc1* deficiency results in impaired NK cell killing. A) Scheme of *invivo* NK cell killing assay. B) Cartoon depicting how WT and β_2_m^-/-^ target cells were labeled for detection by FCM. C and D) Representative FCM plots showing distinct populations of live WT and β_2_m^-/-^ target in poly-I:C-treated WT and *Sostdc1*^-/-^ mice. The left column shows representative FCM plots enumerating the frequencies of WT targets in WT and *Sostdc1*^-/-^ mice. The middle and right column show representative FCM plots enumerating the frequencies of reciprocally labeled β_2_m^-/-^ and WT targets. E) Ratio of targets to show their distribution after 14 hours *in vivo* (Target 1 and Target 2 identified in table below the graph). Each point represents an independent biological replicate. *p<0.05, **p<0.01, Student’s t-test.

### NK cells in Sostdc1^-/-^ mice are impaired in their ability to kill β^2^m-deficient targets

The reduced frequency of splenic mNK with “activating repertoires” that favored activation suggested that NK cell cytotoxicity in *Sostdc1^-/-^* mice would be impaired. To determine if the alterations in NK Ly49 repertoire correlated with NK cell killing ability, we analyzed *Sostdc1*^-/-^ NK cell cytotoxicity with novel FCM-based *in vivo* and *in vitro* killing assays (**Figure 2 and Supplemental Figure 3A-3E**). Beta-2 microglobulin knockout (β_2_m^-/-^) cells express little to no cell surface class I major histocompatibility complex (MHC I) molecules and therefore are sensitive targets for NK cell killing^28^. To test NK cell killing *in vivo*, we pre-activated NK cells in *Sostdc1*^-/-^ and WT control mice with poly(I:C)^29^ **(Figure 2A)** and challenged them with equal numbers of β_2_m^-/-^ and beta-2 microglobulin-sufficient (β_2_m^+/+^) target cells, each labeled with two different fluorochromes (**Figure 2B**). β_2_m^+/+^ target cells from WT and *Sostdc1^-/-^* mice were both included as negative “self” controls (**Figure 2C-2E and data not shown**). After 14 hours of target cell challenge, we quantified the remaining β_2_m^-/-^, WT (β_2_m^+/+^) and *Sostdc1^-/-^* (β_2_m^+/+^) targets by FCM, to determine the frequency of live cells in each target population **(Figure 2A**, **2C and 2D)**, and calculated the ratio of WT (β_2_m^+/+^), *Sostdc1^-/-^* (β_2_m^+/+^), and β_2_m^-/-^ targets in each setting **(Figure 2E)**. An increased proportion of β_2_m^-/-^ targets remained in the *Sostdc1^-/-^* mice compared to WT controls (**Figure 2E**). We confirmed that there was no effect on fluorophore labeling on β_2_m^-/-^ cell target killing with reciprocal labeling of targets of opposing fluorophore (**Figure 2C and 2D**). Thus, these results demonstrated that NK cells in *Sostdc1^-/-^* mice have impaired killing ability (**Figure 2E**).

**Figure 3.**
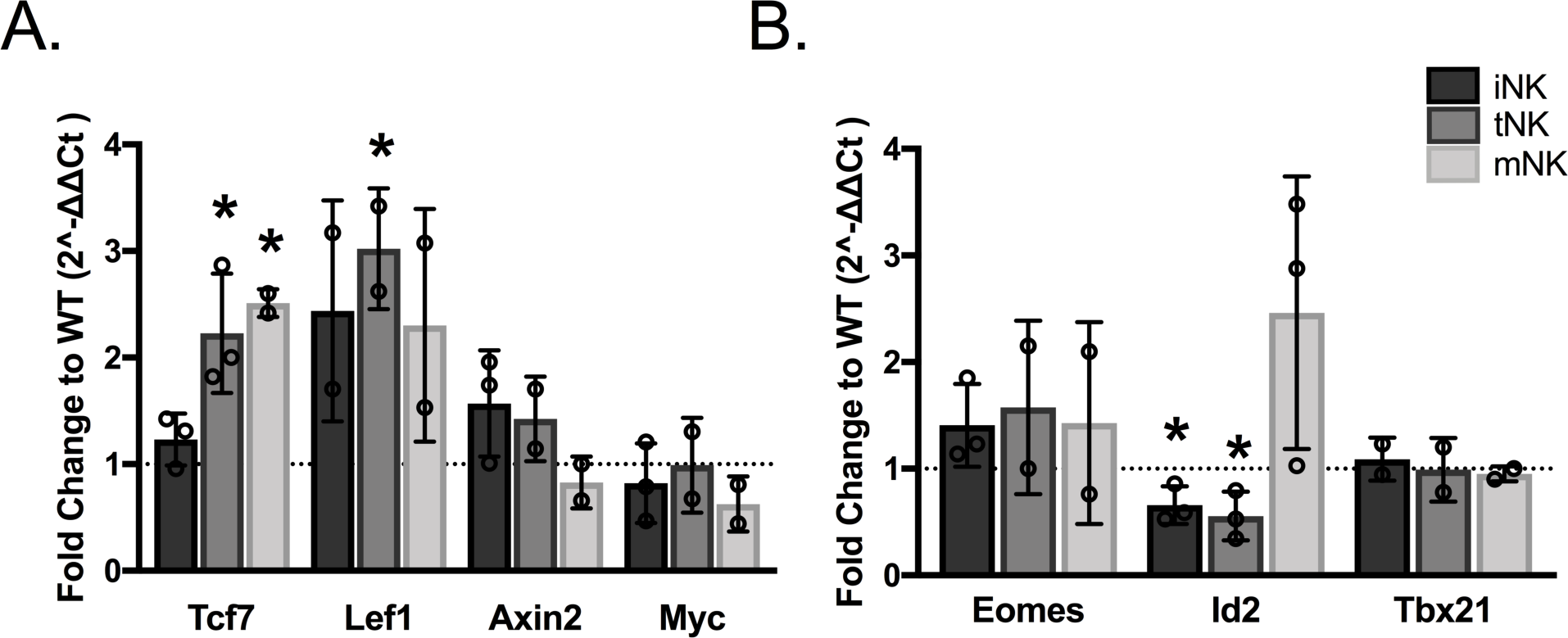
Expression of Wnt pathway and NK cell development genes in *Sostdc1^-/-^* NK cell subsets. A) Gene expression of Wnt pathway genes *Tcf7, Lef1, Axin2,* and *Myc* by quantitative PCR; B) Gene expression of NK cell development genes *Eomes, Id2* and *Tbx21*. Data in A and B are shown relative to WT controls. *p<0.05, Student’s t-test.

This result was confirmed using an *in vitro* NK cell killing assay (**Supplemental Figure 3)** using enriched NK cells from *Sostdc1^-/-^* and WT mice^30^, challenged with fluorescently labeled β_2_m^-/-^ or β_2_m^+/+^ targets for 4 hours, in effector to target (E:T) ratios of 1:1, 2:1, and 4:1 (**Supplemental Figure 3A-3C)**. As expected, WT NK and *Sostdc1^-/-^* NK cells did not lyse β_2_m^+/+^ targets at any E:T ratio (**Supplemental Figure 3D**). However, as shown in **Supplemental Figure 3E**, *Sostdc1^-/-^* NK cells have reduced capacity to lyse β_2_m^-/-^ targets, even at the highest 4:1 E:T ratio, indicating their hyporesponsiveness to β_2_m^-/-^ targets. FCM analysis of *Sostdc1^-/-^* NK cells to produce the cytokine IFNγ after stimulation revealed comparable levels to WT controls **(Supplemental Figure 3F and 3G)**. Surprisingly, activated *Sostdc1^-/-^* NK cells at all developmental stages expressed significantly increased levels of the degranulation marker CD107a **(Supplemental Figure 3H and 3I)**. Taken together, these results suggest that *Sostdc1^-/-^* NK cell cytotoxicity is impaired, despite their ability to produce comparable levels of IFNγ and evidence of elevated accumulation of cytotoxic granules at the cell surface.

### Sostdc1-KO NK cell subsets upregulate Wnt genes Tcf7 and Lef1

Given that Sostdc1 is a known antagonist to canonical Wnt signaling, we hypothesized that expression of canonical Wnt pathway transcription factors would be increased in *Sostdc1^-/-^* NK cell subsets. We purified iNK, tNK, and mNK cells by FCM and analyzed expression of Wnt pathway genes *Tcf7*^3,31^, Lef1^32^, Axin2^33^ and *Myc*^34^, by real-time quantitative polymerase chain reaction (qPCR). Our results showed that relative to WT subsets, *Sostdc1^-/-^* splenic tNK and mNK cells express significantly higher levels of *Tcf7*, and tNK cells also show significantly increased expression of *Lef1* (**Figure 3A**), consistent our hypothesis. Alternatively, we did not observe an increase in *Axin2* and *Myc* expression in any *Sostdc1^-/-^* NK cell subsets (Figure 3A). Together, these results support a role for Wnt signaling by *Tcf7* and *Lef1* in NK cells of *Sostdc1^-/-^* mice.

We also analyzed expression of transcription factors that govern NK cell maturation. T-box family members, Eomesodermin (Eomes) and T-box protein 21 (Tbx21), have been shown to play a crucial role in early immature and mature NK cell maturation^35,36^. Additionally, Eomes-deficient NK cells have reduced Ly49A, Ly49D, Ly49G2, and Ly49H frequencies^35^. Inhibitor of DNA-binding 2 (Id2) is an early TF involved in NK and innate lymphoid cell (ILC) lineage commitment^36,37,38^. *Id2*-deficient mice have fewer mature NK cells and impaired cell killing *in vitro*^39^. Since *Sostdc1^-/-^* mice display a partial maturation block at the tNK cell stage and altered Ly49 receptor frequencies (**Figure 1)**, we hypothesized that we would observe decreased expression of *Eomes, Tbx21, and Id2* at distinct NK cell stages^35–42^. We found *Id2* expression was decreased in iNK and tNK cells in *Sostdc1^-/-^* mice (**Figure 3B**). These results suggest a strong regulation of *Id2* by *Sostdc1* at early NK cell stages, whereas *Eomes* and *Tbx21* are not regulated by *Sostdc1*.

### Sostdc1 in non-hematopoietic stromal cells regulate the maturation of NK cells

To determine if and how *Sostdc1* within specific microenvironmental cell types contributes to the partial block in NK cell maturation and changes in Ly49 repertoires, we performed whole bone marrow transplantation (BMT) experiments. In order to investigate if *Sostdc1* in non-hematopoietic cells influenced NK cell development, we first transplanted whole bone marrow cells from WT(CD45.1^+^/5.1^+^) donors into lethally-irradiated *Sostdc1^-/-^*(CD45.2^+^/5.2^+^) recipients to create WT→KO chimeras. WT(CD45.1^+^/5.1^+^)→WT(CD45.2^+^/5.2^+^) control chimeras were also prepared **(Figure 4A**). Fourteen weeks post-BMT, we analyzed donor-derived NK cell subsets and Ly49 receptor frequencies by FCM. Splenic NK cell numbers were increased (**Figure 4B**), and WT→KO chimeras displayed a partial block between tNK and mNK cell stages in the spleen **(Figure 4C-4E)** and BM (**Supplemental Figure 4C-4E)**, similar to the phenotype that was observed in the non-transplanted *Sostdc1^-/-^* mice (**Figure 1H-1K**). Ly49H^+^ mNK cells were decreased in WT→KO, similar to non-transplanted *Sostdc1^-/-^* mice in the spleen **(Figure 4F and Figure 1N)**. In contrast, WT→KO chimeras contained decreased frequencies of splenic Ly49H^+^ iNK and tNK cells compared to WT→WT controls, a result that was the opposite of the increased frequencies of Ly49H^+^ cells within these NK subsets of non-transplanted *Sostdc1^-/-^* mice **(Figure 4F and Figure 1L-1M)**. In addition, no differences in Ly49G2 and Ly49D subsets were observed between the WT→KO and control chimeras, a result that also differed from the non-transplanted *Sostdc1^-/-^* mice in the spleen **(Figure 4F and Figure 1L-1N)** and BM **(Supplemental Figure 4F)**. Taken together, these results suggested that *Sostdc1* in non-hematopoietic cells controls progression from tNK to mNK stages and NK cellularity, but plays a smaller role in shaping the Ly49 receptor repertoire.

**Figure 4.**
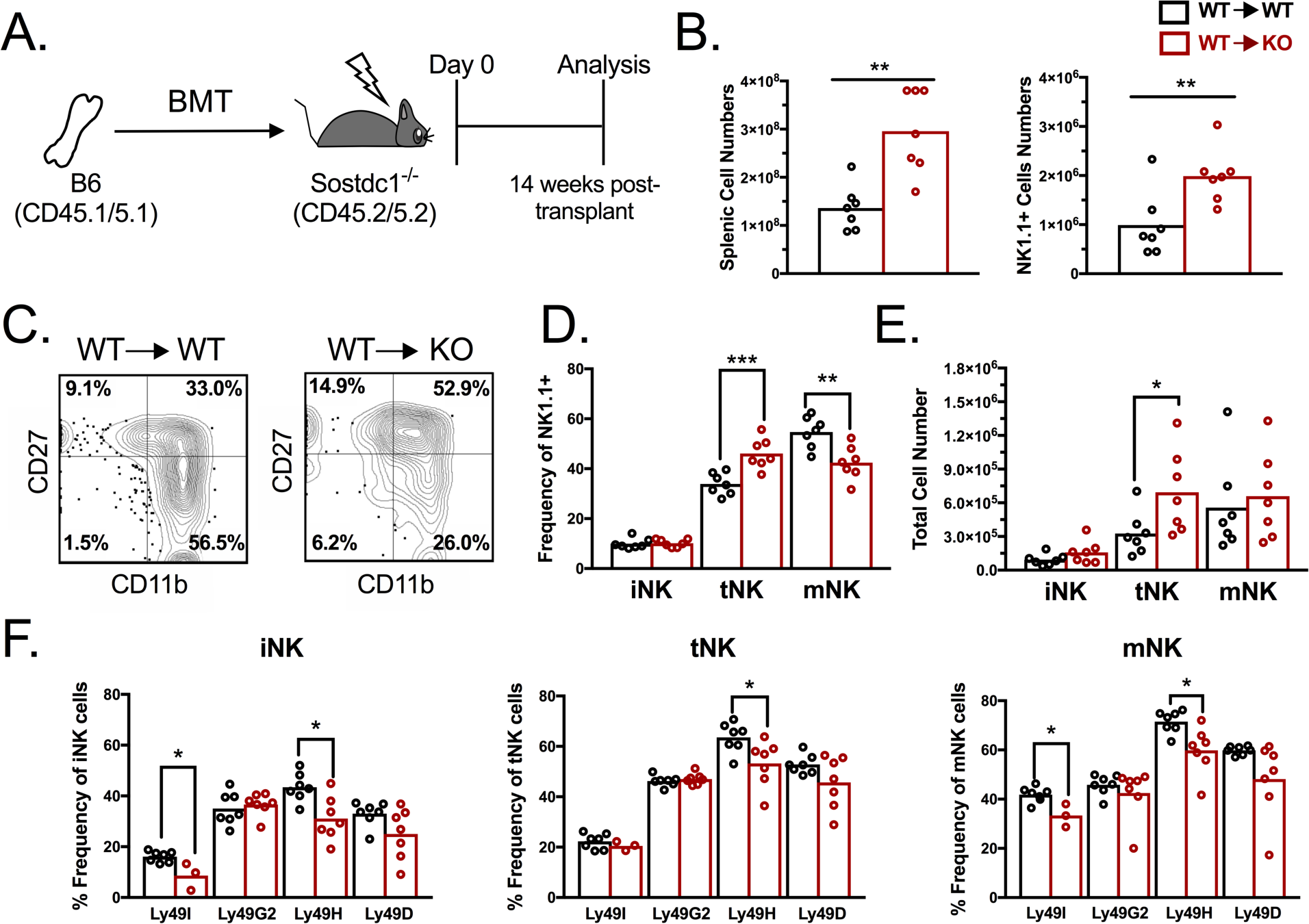
Non-hematopoietic stromal cells regulate the maturation of NK cells. A) Experimental scheme to create WT→*Sostdc1*^-/-^ (WT→KO) bone marrow chimeras. B) Total spleen cellularity (left) and total splenic NK cell numbers in chimeras (right); C) representative FCM plots of NK cell maturation in donor (CD45.1^+^)-derived NK cells in WT→WT and WT→KO chimeras; D) quantification of donor-derived NK cell subset frequencies and E) NK subset cellularity; F) analysis of Ly49 repertoire on donor-derived iNK (left), tNK (center) and mNK (right) cells in WT→WT and WT→KO chimeras. *p<0.05 **p<0.01, ***p<0.001, Student’s t-test.

### Sostdc1 in a hematopoietic cell lineage other than NK cells regulates the Ly49 receptor repertoire

We next prepared reciprocal KO→WT chimeras (whole BM cells from *Sostdc1^-/-^* (CD45.2^+^/5.2^+^) donors transplanted into lethally irradiated WT (CD45.1^+^/5.1^+^) recipient mice) to determine how *Sostdc1^-/-^* NK cells mature and if their Ly49 receptor frequency was changed in a *Sostdc1*-sufficient microenvironment (**Figure 5A**). Remarkably, splenic NK cell numbers **(Figure 5B)** and maturation **(Figure 5C-5E)** were not affected in KO→WT chimeras, in contrast to the non-transplanted *Sostdc1^-/-^* mice and the WT→KO chimeras **(Figure 1H-1I and Figure 4B-4E, respectively)**. However, the BM analysis showed an increase in NK cell numbers **(Supplemental Figure 4H)** and a similar tNK cell accumulation as observed in the non-transplanted *Sostdc1^-/-^* mice (**Figure 1H-1K** and **Supplemental Figure 4K).** Furthermore, analysis of donor-derived Ly49-expressing NK cell subsets in the spleens in KO→WT chimeras demonstrated some similar patterns as non-transplanted *Sostdc1^-/-^* mice, such as the increase in frequencies of Ly49H^+^ and Ly49D^+^ tNK cells and a decrease in the frequencies of Ly49G2^+^ iNK cells, and decreased frequencies of Ly49G2^+^ and Ly49I^+^ mNK cells **(Figure 5F and Figure 1L-1N).** However, higher frequencies of mNK cells expressing Ly49H and Ly49D were observed in the KO→WT spleens, whereas these populations were decreased in non-transplanted *Sostdc1^-/-^* mice **(Figure 5F and Figure 1L-1N)**.

**Figure 5.**
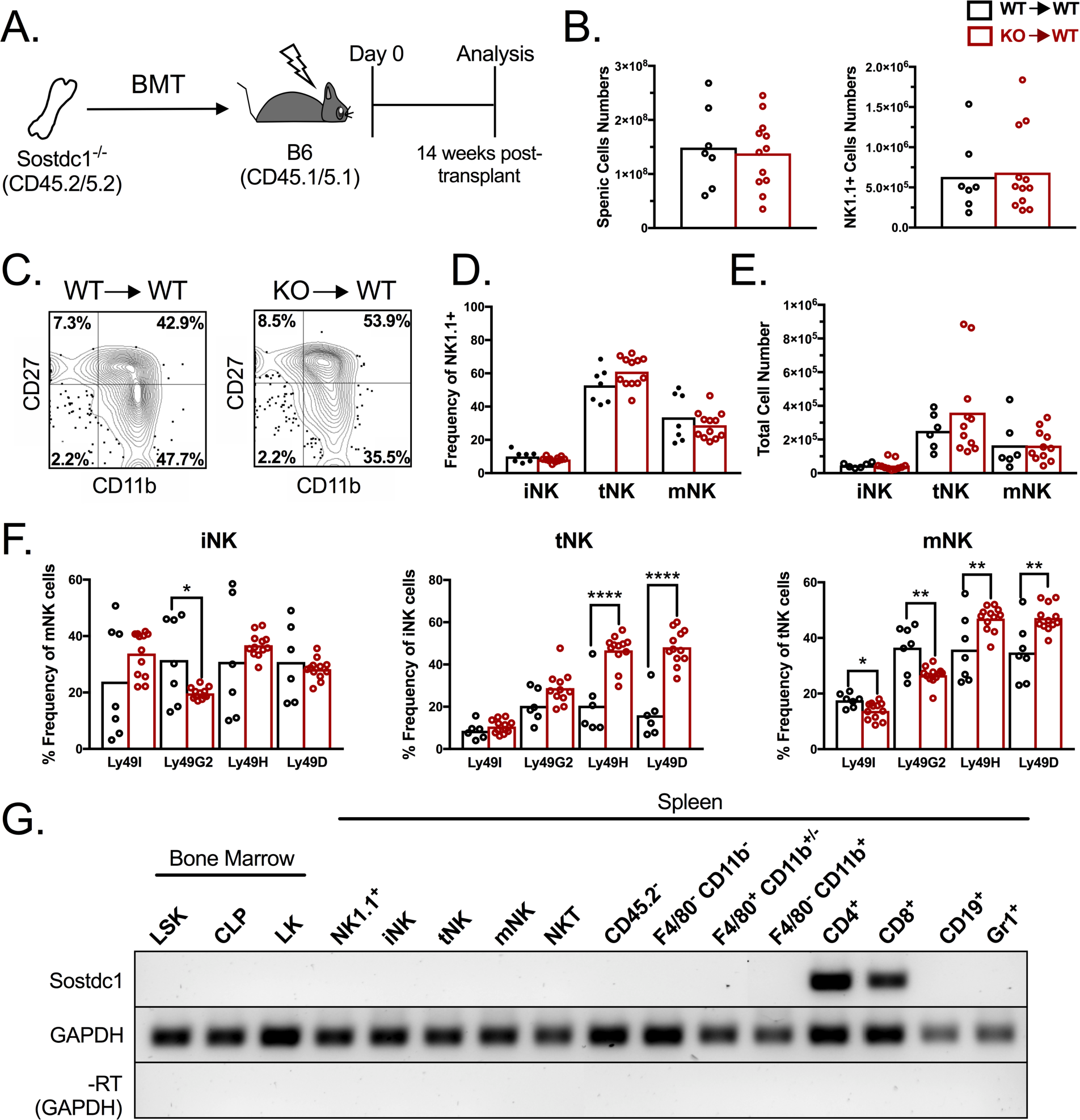
Sostdc1 in hematopoietic non-NK cells regulates the Ly49 NK receptor repertoire. A) Experimental scheme to create *Sostdc1*^-/-^→WT (KO→WT) bone marrow chimeras. B) total cellularity in spleen (left) and total donor (CD45.2^+^)-derived NK cell numbers (right) in chimeras; C) representative FCM plots of splenic NK cell maturation of donor-derived NK cells in WT→WT and KO→WT chimeras; D) quantification of donor NK cell subset frequencies and E) NK cellularity in the spleen of chimeras; F) analysis of Ly49 repertoire on donor-derived iNK (left), tNK (center) and mNK (right) cells in WT→WT and KO→WT spleens. G) *Sostdc1* expression in hematopoietic cell lineages by RT-PCR. *p<0.05 **p<0.01, ***p<0.001, Student’s t-test.

The Ly49 frequency patterns observed in the KO→WT chimeras strongly suggested that *Sostdc1* in NK cells regulated the Ly49 repertoire in a cell-intrinsic fashion. To confirm this, we examined *Sostdc1* expression in sorted WT iNK, tNK, and mNK cells by qPCR. Surprisingly, iNK, tNK, and mNK cells do not express *Sostdc1* (**Figure 5G**). To determine alternative possible hematopoietic sources of *Sostdc1*, we examined other populations, such as Lineage^-^Sca1^+^Kit^+^ (LSK), Lineage^-^Kit^+^Sca1^-^ (LK), and common lymphoid progenitors (CLP), NK T cells, macrophages, B cells and granulocytes, which were all negative for *Sostdc1* **(Figure 5G)**. Only CD4^+^ and CD8^+^ T cells displayed high levels of *Sostdc1* expression (**Figure 5G**). This expression pattern was confirmed in CD4 and CD8 T cells from *Sostdc1^-/^*^-^ mice using PCR for LacZ^26^ **(data not shown)**. Collectively, these results indicate that *Sostdc1* does not regulate splenic NK cell development in a NK cell-intrinsic manner, and identifies *Sostdc1*-positive T cells as putative NK niche cells that may contribute to shaping of the Ly49 repertoire.

## Discussion

We have uncovered novel roles of the *Sostdc1* gene in NK cell maturation and function through two distinct mechanisms. Our working model is illustrated in **Figure 6**. Our data supports that Sostdc1 from two distinct sources, non-hematopoietic stromal cells, and hematopoietic cells (in particular, CD4^+^ and CD8^+^ T lymphocytes) regulate NK cell maturation versus Ly49 receptor expression and frequencies, respectively, in somewhat independent manners, and that this occurs through the control of Wnt signaling activation. Our developmental and functional NK cytotoxicity assay results lead us to conclude several NK “niche cell” populations exist that require Sostdc1 expression in order to produce a healthy NK cell repertoire that can distinguish between self and non-self.

**Figure 6.**
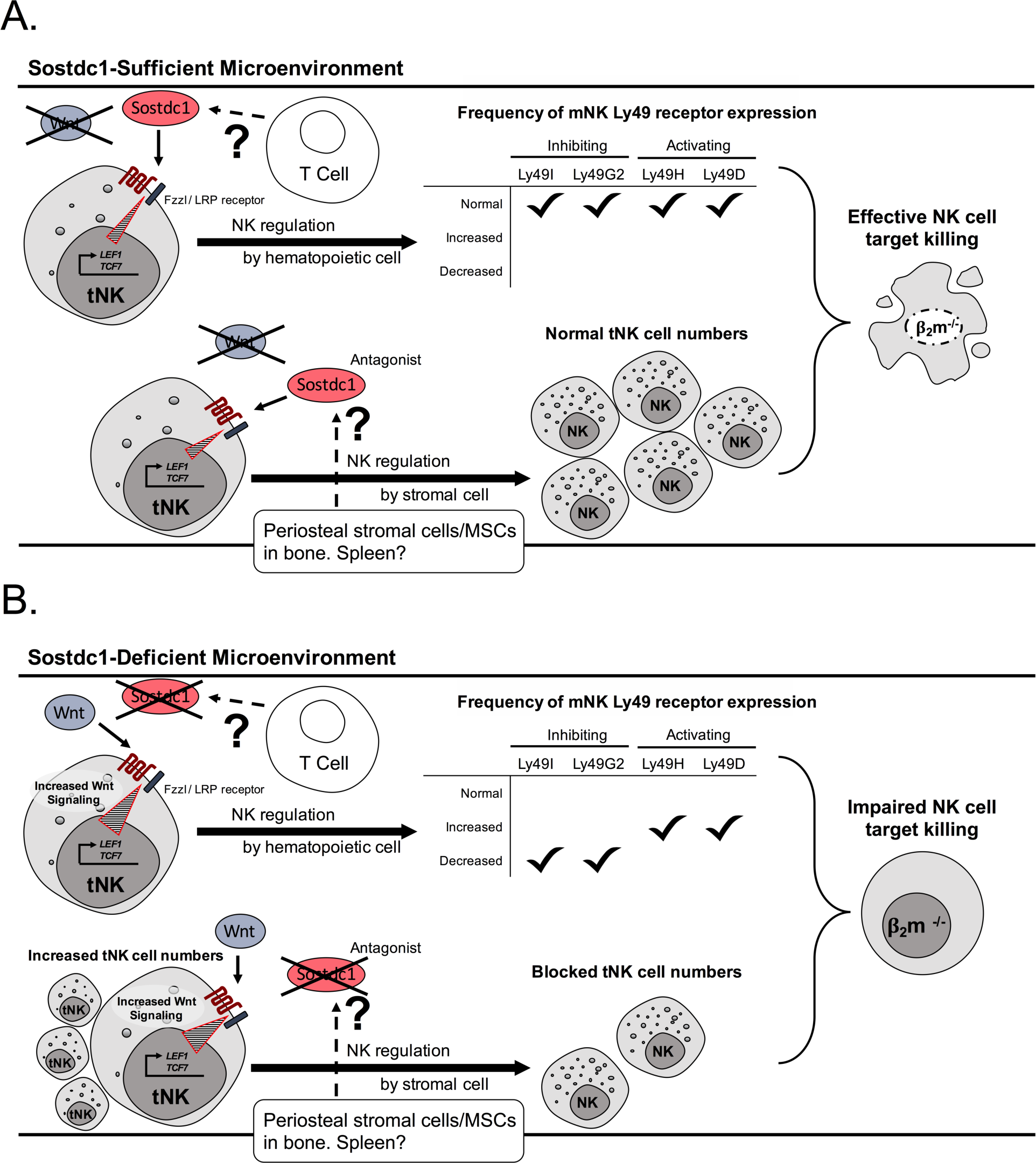
Working model of Sostdc1’s role in NK cell maturation and cytotoxicity. A)In a *Sostdc1*-sufficient (wild-type) microenvironment, *Sostdc1* is expressed by non-hematopoietic stromal cells in the bone that antagonizes Wnt signaling in NK cells, resulting in baseline levels of *Tcf7* and *Lef1*, and regulates NK cell numbers. *Sostdc1* expressed by T cells also influences Wnt signaling genes, but distinctly controls the distribution of the Ly49 repertoire. Collectively, Sostdc1 is required in the microenvironment for development of NK cells with the ability to effectively recognize, be primed for activation and lyse MHC-I deficient targets. B) In the absence of *Sostdc1* in bone stromal cells, splenic stromal cells or T cells, *Tcf7* and *Lef1* expression is increased as a result of overstimulated Wnt signaling, which adversely affects NK cell numbers. Loss of *Sostdc1* in T cells might also dysregulate the distribution of Ly49s amongst NK cell subsets. Collectively, loss of *Sostdc1* in stromal cells and T cells acts cell-extrinsically on NK cells, producing NK cells that are hyporesponsive to MHC-I deficient targets.

Perhaps the most surprising finding from our studies is that *Sostdc1* is not expressed in NK cells themselves, and that Ly49 receptor expression in the spleen is possibly controlled in a cell-extrinsic manner by CD4^+^ and CD8^+^ T cells. To our knowledge, there is no previous demonstration of NK and T cell crosstalk that shapes the NK Ly49 receptor profile. Recent studies have defined *Sostdc1* as a marker of relatively rare memory PD-1^+^ CD4^+^ T cells^43,44^ and T follicular helper (T_FH_) cells found in the Peyer’s Patches and peripheral lymph nodes^10^, but only cite Sostdc1’s role on humoral immunity. We observed high expression of *Sostdc1* in bulk sorted splenic CD4^+^ and CD8^+^ cells from the spleen, and we assume that most of the *Sostdc1* expression is coming from PD1^+^ CD4^+^ T cells and T_FH_ cell subsets, based on these published studies. We have not observed any obvious block in T cell development in the *Sostdc1*^-/-^ mice (data not shown). Undeniably, further experimentation is required to definitively connect the role of these T cell subsets in NK cell regulation, and to determine whether the T cells are regulating Ly49 receptor expression via directly binding to NK cells, or mediating their effects indirectly by secreting Sostdc1 in a paracrine fashion.

Wnt signaling has been well studied in the framework of hematopoietic stem cells (HSCs) to help promote proliferation, differentiation, and homeostasis^45,46^. It is now evident that canonical Wnt signaling plays a crucial role in the regulation of many immune cells^45^. Based on the antagonist role of Sostdc1 on Wnt signaling and our discovery of increased *Tcf7* and *Lef1* expression in NK cells from *Sostdc1^-/-^* mice, we conclude that canonical Wnt signaling plays a crucial role in NK cell development and function, particularly at the tNK cell stage. In the absence of *Sostdc1*, tNK cells are partially blocked in their maturation, express dysregulated Ly49 frequencies in the BM and spleen, and appear to be more reliant on Wnt activation. Our observations that one of the coactivators of canonical Wnt signaling, Tcf7, was significantly upregulated in tNK and mNK cells, and the secondary coactivator, Lef1, was significantly upregulated in the mNK cell stage, suggest that a critical period exists in which tNK cells require downregulation of Tcf7 and Lef1 in order to progress to the mNK cell stage. Our results and interpretation are consistent with a recent study that downregulation of Tcf1 (encoded by *Tcf7*) is required for full NK cell maturation and cytotoxicity^3^.

Based on our current findings, we cannot rule out whether the impaired killing of β_2_m^-/-^ targets by NK cells from *Sostdc1*^-/-^ mice is due to insufficient numbers of mNK cells, inefficient execution of the perforin and granzyme pathways, the dysregulation of Ly49 receptor frequencies amongst NK cells, or a combination of all of these possible mechanisms. NK cells at tNK and mNK stages express genes involved in cytotoxic function^15,11,12^. It has been shown that Ly49 receptor expression is required for NK cell cytoxicity^47^, which is consistent with our observations that the splenic mNK cells in the *Sostdc1*^-/-^ mice contain decreased frequencies of all Ly49-expressing subsets and the killing ability of *Sostdc1^-/-^* NK cells towards β_2_m^-/-^ targets is poor. Because *Sostdc1*^-/-^ mice express increased proportions of tNK cells with an “activating” repertoire^48^ and high CD107a levels, we would have expected enhanced target cell killing by the *Sostdc1*^-/-^ NK cells, but we observed them to be hyporesponsive. Taken together, our results suggest that NK cell cytotoxicity is universally disabled in the absence of *Sostdc1*, but is caused by distinct mechanisms in tNK and mNK cells. Further experiments, in which the killing ability of purified tNK and mNK cells from *Sostdc1*^-/-^ mice is specifically examined, are necessary to definitely demonstrate this. Additional work is also needed to dissect the specific roles of non-hematopoietic and hematopoietic cells on NK cell cytotoxicity. Understanding the details of the basic biology underlying the development and regulation of NK cell cytotoxicity and how these processes are quantitatively integrated could be applied to manipulate these processes in a controlled fashion in order to produce specific numbers of NK cells with enhanced killing ability, and perhaps impact the production of NK cell-based cancer immunotherapies^1–6^.

## Materials and Methods

### Mice

*Sostdc1*^-/-^ mice have been described^22,26^, and both bred and transferred from Lawrence Livermore National Laboratories (LLNL) to UC Merced to begin an independent breeding colony. C57B6/J (CD45.2^+^/5.2^+^) and B6.SJL-Ptprca Pepcb/BoyJ (CD45.1+/5.1+) and B6.129P2-B2mtm1Unc/J (β_2_m^-/-^) mice were obtained from Jackson Laboratories. Mice of 28-38 weeks of age and of both sexes were used. No differences between sexes nor mice from LLNL and UC Merced colonies have been observed. All mice were housed in conventional housing with autoclaved feed. Mice were euthanized by carbon dioxide asphyxiation followed by cervical dislocation. All animal procedures were approved by the UC Merced and LLNL Institutional Animal Care and Use Committees.

### Flow cytometry

Isolation of spleen and bone marrow cells were performed and stained for flow cytometry as described^26^. Antibodies against CD161 or NK1.1 (PK136), CD11b (M1/70), CD27 (LG.3A10), CD19 (6D5), CD3 (2C11), Gr1-Ly6C/G (Gr1), Ly49G2 (4D11), Ly49I (YL1-90), Ly49H (3D10), Ly49D (eBio4E5), CD45.2 (104), CD45.1 (A20), CD45 (30-F11), CD4 (GK1.5), CD8 (2.43), Ter119 (TER119), CD107a (1D4B), Rat IgG2a κ Isotype Control (RTK2758), IFNγ (XMG1.2), Rat IgG1𝜅𝜅 Isotype control (RTK2071), and BUV395 Streptavidin, eFluor 780 Fixable Viability Dye, eFluor 506 Fixable Viability Dye were purchased from eBioscience, BioLegend, Miltenyi Biotec, and BD Biosciences. Staining of all cells included a pre-incubation step with unconjugated anti-CD16/32 (clone 2.4G2 or clone 93) mAb to prevent nonspecific binding of mAbs to FcγR. For extracellular staining, the cells were washed and incubated with a panel of mAbs for 15-20 minutes at 4C or on ice, and then washed again. Intracellular staining was performed using the BD Cytofix/Cytoperm Fixation/Permeabilization Kit (BD Biosciences) per the manufacturer’s instructions. To purify NK cells and NK cell subsets, enrichment of NK cells was first achieved by staining with biotinylated anti-“lineage” cocktail (anti-CD3, CD4, CD8, CD19, Gr1, and Ter119) followed by magnetic separation using EasySep Positive Selection kit (Stem Cell Technologies), After enrichment, cells were stained with streptavidin-FITC, anti-CD27, and additional anti-CD3, CD19, Gr1, CD45, NK1.1, and CD11b. Lineage-negative CD45^+^ NK1.1^+^ iNKs, tNKs and mNKs were sorted on the FACS Aria II (Becton Dickinson). Flow cytometric data was acquired on the BD LSR II or FACS Aria II cell sorter (Becton Dickinson). The data was analyzed using Flowjo version 7.6 or 10 (Tree Star, Inc.)

### In vivo NK cell killing assay

*Sostdc1*^-/-^ and B6 control mice received 200 μg of polyinosinic-polycytidylic acid (Poly(I:C)) (Sigma-Aldrich) via injection into the intraperitoneal cavity. Thirty-six hours later, splenic cells from aged and sex-matched β_2_m^-/-^ and β_2_m^+/+^ (WT or *Sostdc1^-/-^)* mice were harvested and processed to a single cell suspension in media (Medium 199, 2% FCS, 2mM L-glutamine, 100U/ml penicillin, 100 μg/ml streptomycin, 25 mM HEPES) and counted using a hemocytometer. β_2_m^+/+^ (WT or *Sostdc1^-/-^*) and β_2_m^-/-^ control target cells were stained with anti-CD45 conjugated to either APC or PE in media for 20 min on ice. 5×10^6^ β_2_m^-/-^ stained splenic cells were mixed with 5×10^6^ β_2_m^+/+^ stained control cells at a 50:50 ratio, thus providing a method to track each target cell type by flow cytometry. Stained cell target cell mixtures were injected intravenously by retro-orbital injection. Fourteen hours later, spleens from *Sostdc1*^-/-^ and WT recipients were then harvested and processed for flow cytometry. NK cell lysis of targets was determined by flow cytometry (FCM) and calculated by the ratio of live β_2_m^+/+^ (WT), β_2_m^+/+^ (*Sostdc1^-/-^*) and β_2_m^-/-^ targets over WT control cells in the same mouse.

### Gene expression analysis by quantitative PCR

Cells were pelleted and resuspended in RNeasy Lysis Buffer with 2-mercaptoethanol (Qiagen). Total RNA was purified using Qiagen RNeasy Mini Kit (Qiagen) according to manufacturer’s protocol. RNA concentration and purity was analyzed using a NanoDrop ND-1000 Spectrophotometer (Thermo Fisher Scientific). iScript cDNA Synthesis Kit was used (Bio-Rad Laboratories) according to the manufacturer’s protocol. Real-time quantitative PCR performed using the iTaq Universal SYBR Green Supermix kit (Bio-Rad Laboratories) and ran on a Stratagene Mx3000P thermocycler (Thermo Fisher Scientific) using the following conditions: 1 cycle at 9°C for 30 s, followed by 40 cycles of 95°C for 5 s and 60°C for 30 s, and a final cycle at 95°C for 1 min, 55°C for 30 s, and 95°C for 30 s to end the run. The PCR products were visualized on a 2% agarose gel and imaged under UV light using a ChemiDoc (Bio-Rad Laboratories) with SYBR Safe (Invitrogen) stain. The list of genes and their primer sequences are listed in Supplementary Table 1.

### Bone marrow chimeras

Whole BM cells were aseptically isolated from B6 (CD45.1 or CD45.2) wild type (WT) or *Sostdc1*^-/-^ (CD45.2) mice and 5×10^6^ cells were transferred via retro-orbital injection into lethally (10 Gy) irradiated recipients 4 hours after irradiation using a cesium irradiator. Mice were given neomycin-containing drinking water for 2 weeks post-transfer. Chimeras were analyzed 14 weeks post-transplant.

## Acknowledgments

This work was supported by University of California (UC), Merced faculty research funding, University of California Cancer Research Coordinating Committee grant and Halcyon-Dixon Trust award to JOM, and UC Graduate Student Fellowships to AM. GGL works under the auspices of the U.S. Department of Energy by Lawrence Livermore National Laboratory under Contract DE-AC52-07NA27344. Authors’ roles: GGL and DM provided the *Sostdc1^-/-^* mice. Study design: AM and JOM. Study conduct: AM, SE, EL, JA, JOM. Data collection: AM, SE, EL, JA. Data analysis: AM, SE, EL, JA, JOM. Data interpretation: AM and JOM. Drafting manuscript: AM and JOM. Revising manuscript content: AM and JOM. Approving final version of manuscript: JOM. JOM takes responsibility for the integrity of the data analysis. The authors thank the staff of the Department of Animal Research Services and the Flow Cytometry Core of the Stem Cell Instrumentation Foundry at UC Merced for excellent animal care and technical support, and for Dr. Anna Beaudin and Dr. Marcos E. García-Ojeda for their comments on the manuscript.

